## Supplementary figures for "A whole-organism landscape of X-inactivation in humans"

### Supplementary Figure 1

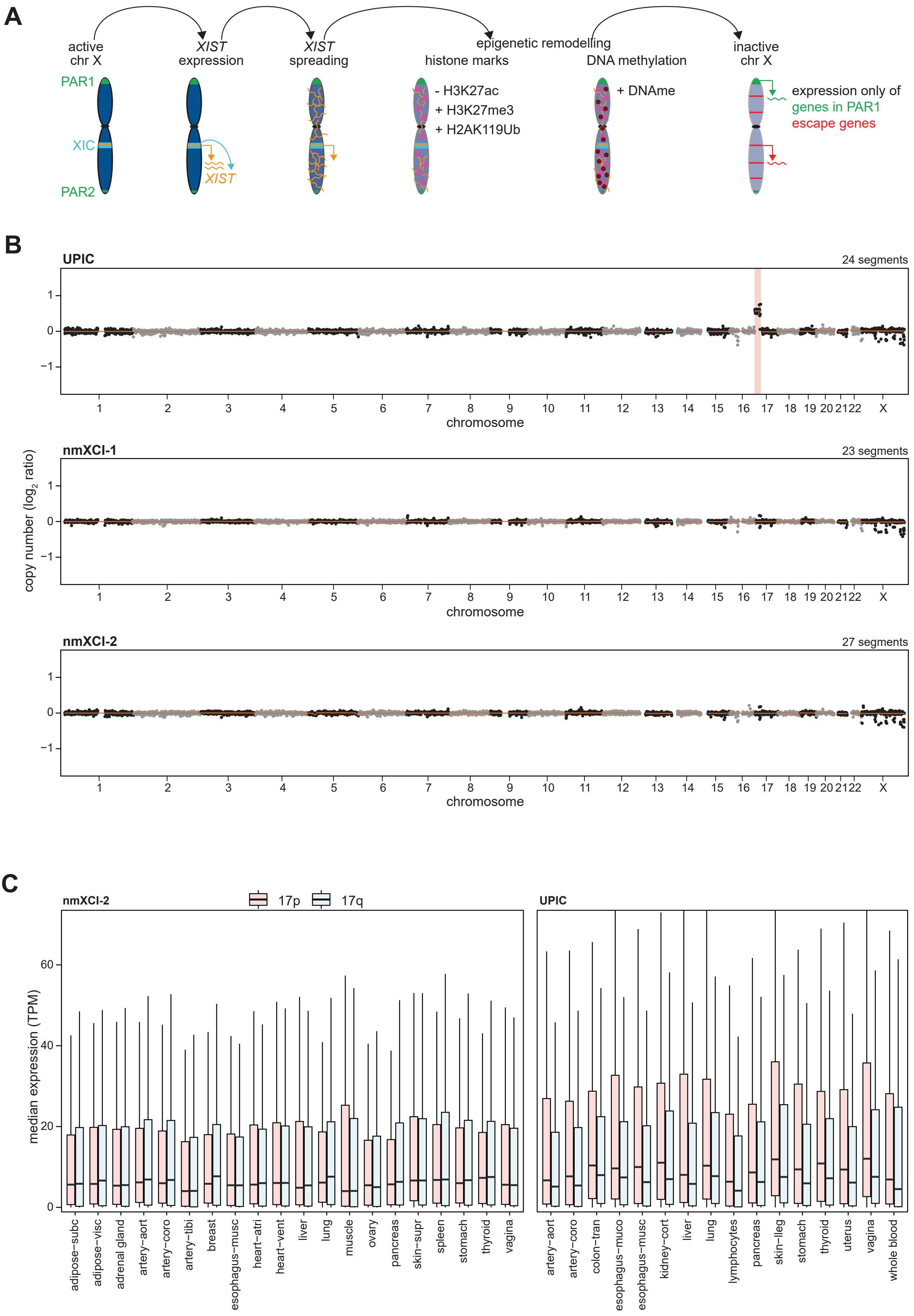

### Supplementary Figure 2

**A**

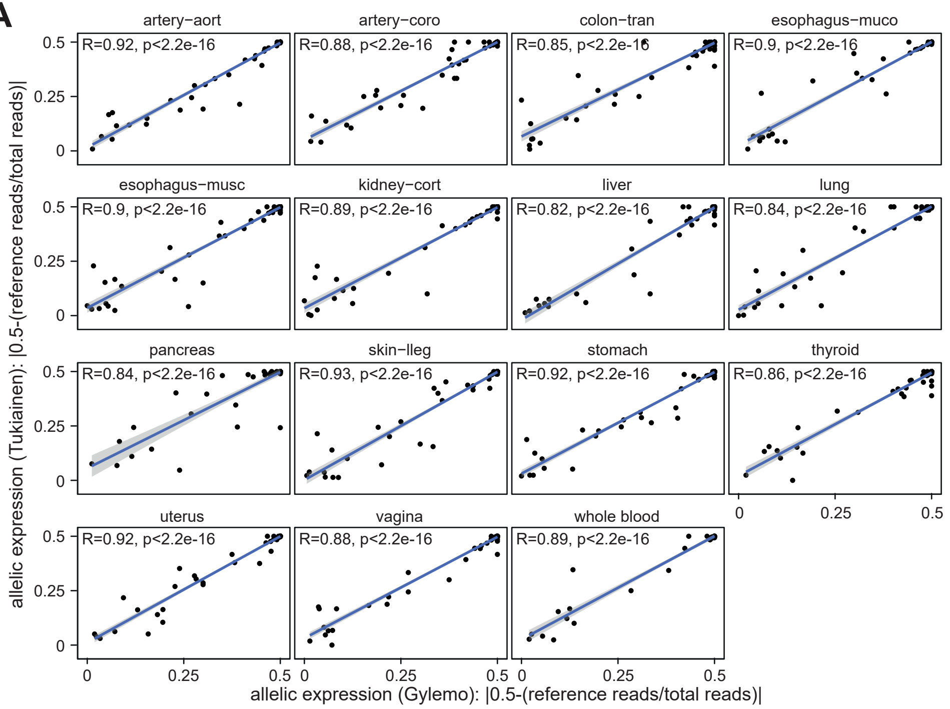


**B**

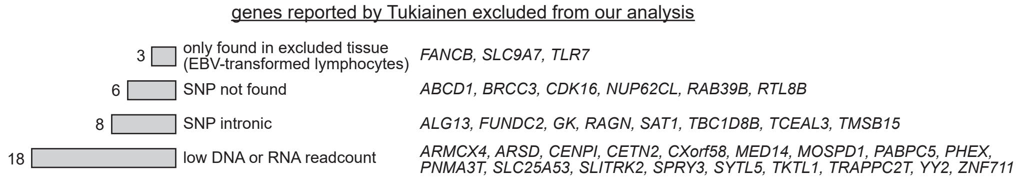

### Supplementary Figure 3

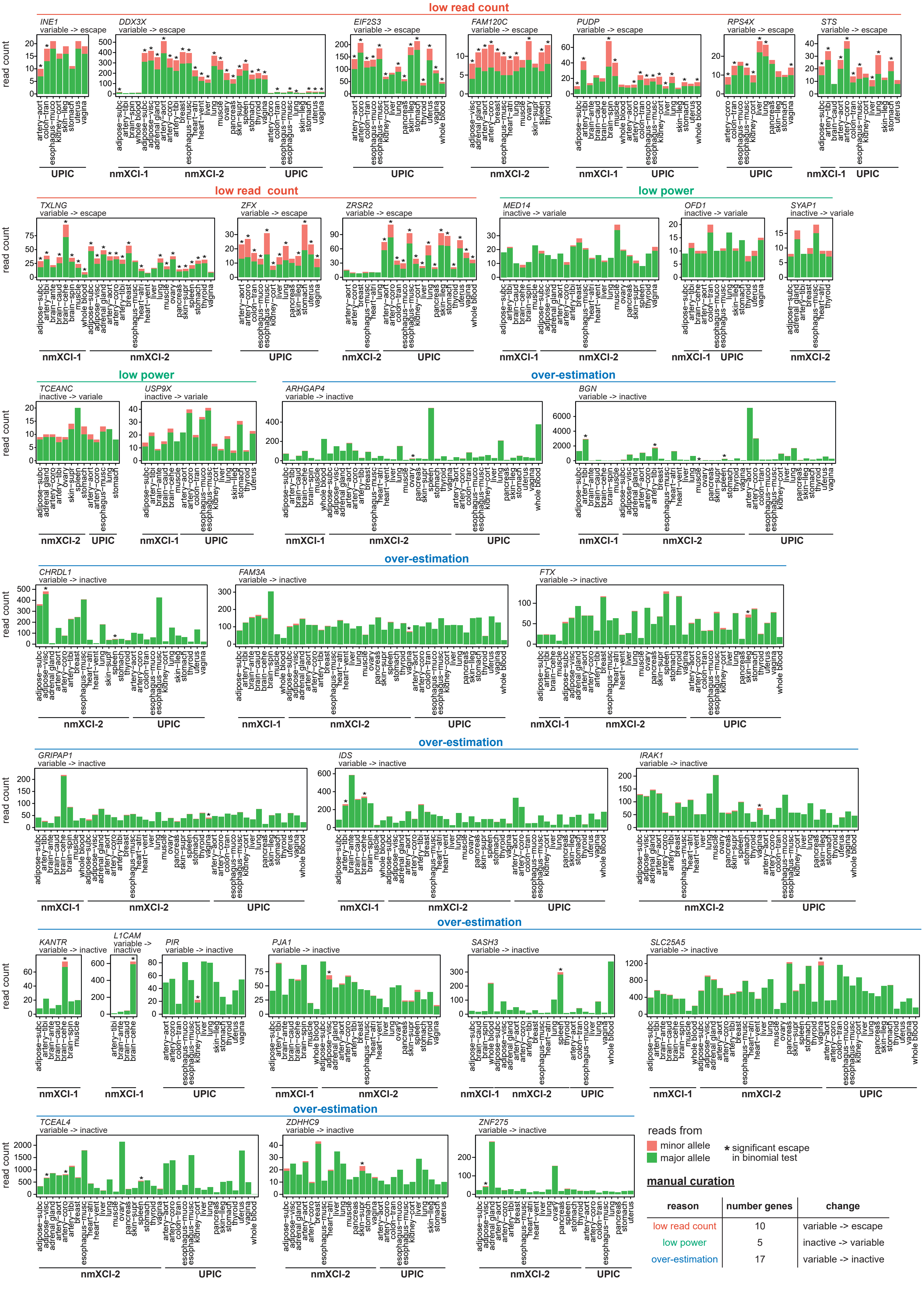

### Supplementary Figure 4

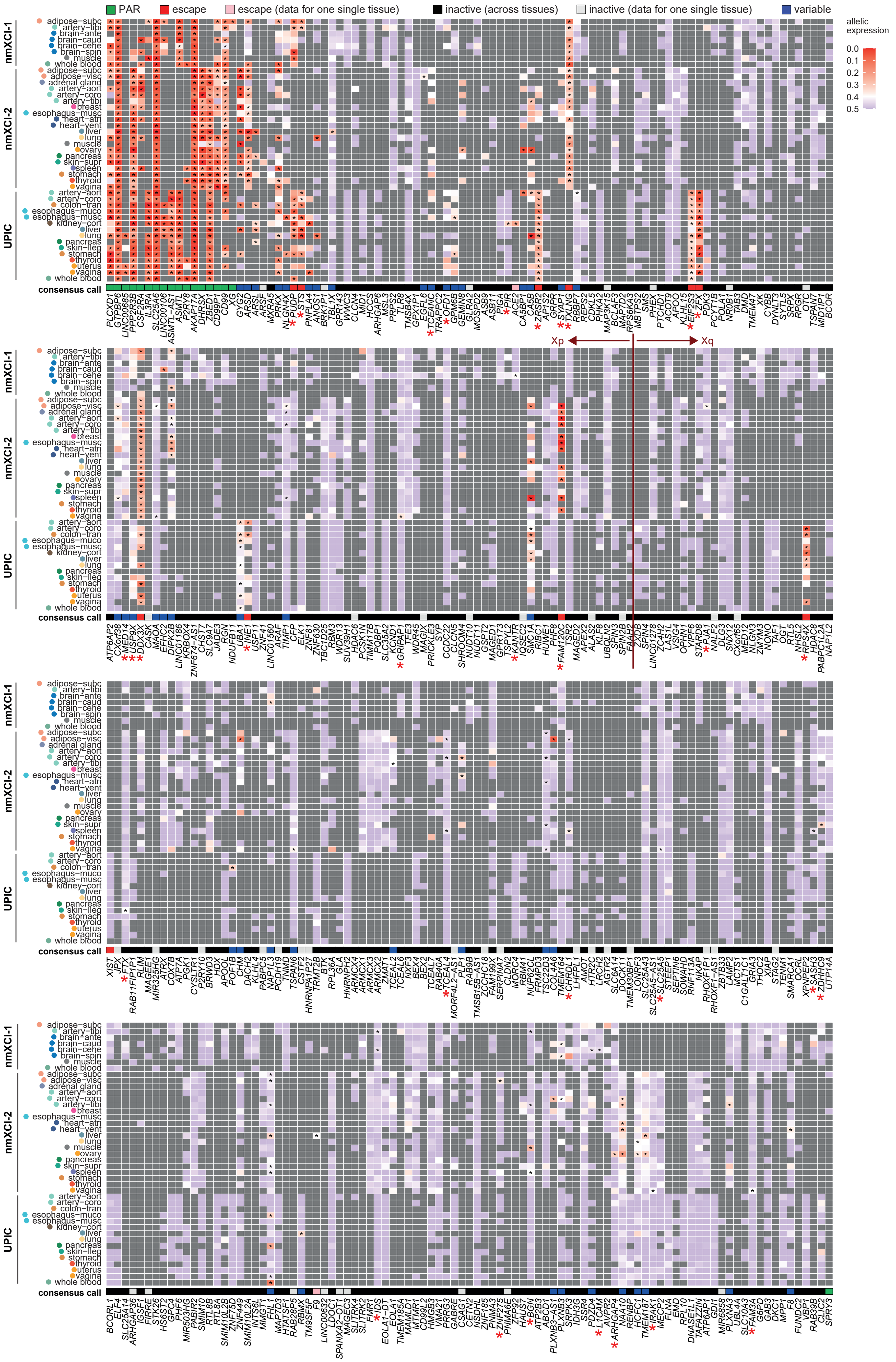

### Supplementary Figure 5

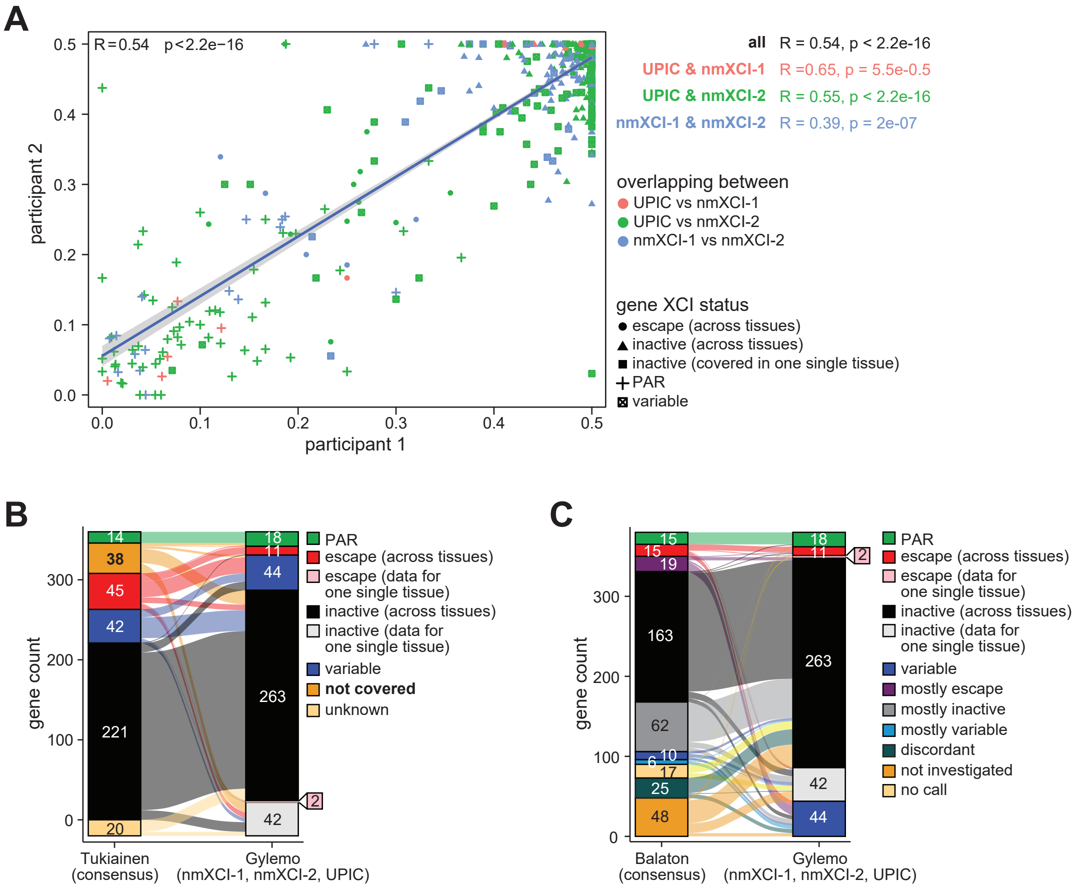
